# Tacc3 modulates microtubule network dynamicity and focal adhesion remodeling to affect cranial neural crest cell migration in *Xenopus laevis*

**DOI:** 10.1101/2021.02.08.430297

**Authors:** Elizabeth A. Bearce, Benjamin Pratt, Erin Rutherford, Leslie Carandang, Laura Anne Lowery

## Abstract

Coordinated cell migration is critical during embryogenesis, as cells must leave their point of origin, navigate a complex barrage of signals, and accurately position themselves to facilitate correct tissue and organ formation. The cell motility process relies on dynamic interactions of the F-actin and microtubule (MT) cytoskeletons. Our work focuses on how one MT plus-end regulator, Transforming Acidic Coiled-Coil 3 (Tacc3), can impact migration of cranial neural crest cells in *Xenopus laevis*. We previously demonstrated that *tacc3* expression is expressed in cranial neural crest cells, and that Tacc3 can function as a MT plus-end tracking protein to regulate MT growth velocities. Here, we demonstrate that manipulation of Tacc3 protein levels is sufficient to alter cranial neural crest cell velocity *in vitro*. Tacc3 overexpression drives increased single-cell migration velocities, while Tacc3 KD results in reduced cell velocity and defective explant dispersion. We also show that Tacc3 can have spatially-enhanced effects on MT plus-end growth velocities as well as effects on focal adhesion remodeling. Together, we demonstrate that Tacc3 can facilitate neural crest cell motility through spatially-enhanced cytoskeletal remodeling, which may underlie the enhanced metastatic potential of Tacc3-overexpressing tumor cells.

## Introduction

Properly-coordinated cell migration is a pivotal aspect of embryogenesis and requires integration of dynamic interactions between F-actin, actomyosin forces, and focal adhesions (FAs) (Gardel, Schneider et al. 2010; Blanchoin, Boujemaa-Paterski et al. 2014; Puleo, Parker et al. 2019; Schaks, Giannone et al. 2019). Additionally, microtubules (MTs) and their associated proteins (MAPs) play integral roles during directed cell motility, by way of modulating FA disassembly, contributing to cell polarity, and facilitating delivery of cell components to the leading edge (Etienne-Manneville 2013; Bouchet and Akhmanova 2017; Garcin and Straube 2019). One family of MAPs, the MT plus-end tracking proteins (+TIPs), is especially well-situated to influence leading edge dynamics (Galjart 2010). MT plus-ends are largely biased towards the leading edge of motile cells, as they are in a prime location to sense changes in the extracellular environment, affect MT network stability, and mediate crosstalk between MTs, actin, and FAs (Wittmann and Waterman-Storer 2005; Kumar, Lyle et al. 2009; Gierke and Wittmann 2012; Stehbens, Paszek et al. 2014; Bearce, Erdogan et al. 2015; Efimova, Yang et al. 2020; Juanes, Fees et al. 2020; Sanchez-Huertas, Bonhomme et al. 2020).

One promising +TIP for regulating cytoskeletal interactions to promote cell migration is Tacc3, a member of the Transforming acidic coiled-coil (TACC) family. While Tacc3 is best known for regulating MT stabilization at the mitotic spindle (Gergely, Kidd et al. 2000; Ding, Huang et al. 2017; Burgess, Mukherjee et al. 2018; Ryan, Shelford et al. 2020), Tacc3 also has critical functions in interphase cells. For example, we previously showed that Tacc3 functions as a +TIP in interphase embryonic cells and promotes axon extension and guidance cue response (Nwagbara, Faris et al. 2014; Erdogan, Cammarata et al. 2017; Erdogan, St Clair et al. 2020). Moreover, we observed that Tacc3 expression is enriched in the cranial neural crest (CNC) cells in *Xenopus* (Mills, Bearce et al. 2019), while others have shown TACC3 to be expressed in murine embryonic tissues as well (Aitola, Sadek et al. 2003). Murine embryos that were TACC3-depleted showed severe facial clefts, a well-known indicator of deficits in CNC cell motility (Piekorz, Hoffmeyer et al. 2002). Additionally, TACC3 is frequently dysregulated in multiple types of cancer, and this is increasingly correlated with metastatic potential and poor survival prognoses (Lauffart, Vaughan et al. 2005; L’Espérance, Popa et al. 2006; Yim, Tong et al. 2009; Ha, Kim et al. 2013; Ha, Park et al. 2013; Li, Ye et al. 2017; Matsuda, Miyoshi et al. 2018; Song, Liu et al. 2018; Qie, Wang et al. 2020). While TACC3 may play a role in promoting cell proliferation in cancer, there is also evidence that TACC3 drives increased cell invasiveness and metastasis (Ha, Kim et al. 2013; Zhu, Cao et al. 2016; Li, Ye et al. 2017; Guo and Liu 2018; Zhao, He et al. 2018). However, the specific mechanisms by which TACC3 regulates cell migration are still unclear.

Here, we sought to examine whether Tacc3 plays a direct role in cytoskeletal regulation during CNC cell motility. We demonstrate that Tacc3 manipulation can drive changes in mean cell migration velocity and alter cell-to-cell ensemble dispersion. MT dynamics data suggest these impacts may stem from the ability of Tacc3 to enhance MT network plasticity. We also demonstrate that Tacc3 can affect FA remodeling. We predict that, during CNC cell migration, these effects can drive enhanced cytoskeletal repolarization following cell-cell contact, one of the predominant mediators of developmental cell motility. We suggest that maintaining a plastic MT network that can react resiliently during random migration may convey advantages to the cancer during metastasis.

## Results and Discussion

### Tacc3 manipulation impacts cell velocity and dispersion dynamics of CNC cells

After CNC cells are specified along the anterior neural tube, they undergo an epithelial-to-mesenchymal transition, delaminating to migrate ventrally into streams called the pharyngeal arches (Fukiishi and Morriss-Kay 1992; Cordero, Brugmann et al. 2011). We previously demonstrated that Tacc3 transcripts are expressed in motile CNC cells of the pharyngeal arches in *Xenopus laevis* (Mills, Bearce et al. 2019). Moreover, we found that whole-embryo Tacc3 depletion resulted in shorter CNC cell streams (**Figure S1**), potentially indicating that Tacc3 reduction could negatively impact CNC cell motility. However, shortened CNC streams could also be explained by a reduction in total number of cells. To distinguish between these possibilities, we dissected out CNC cells from control and Tacc3-depleted *Xenopus laevis* embryos that were co-injected with a nuclear marker and allowed to migrate freely on fibronectin-coated coverslips (**Figure 1**). Cell nuclei were segmented and tracked to ascertain cell velocity and directionality (**Figure 1A-F**), using automated particle tracking (Tinevez, Perry et al. 2017).

**Figure 1:**
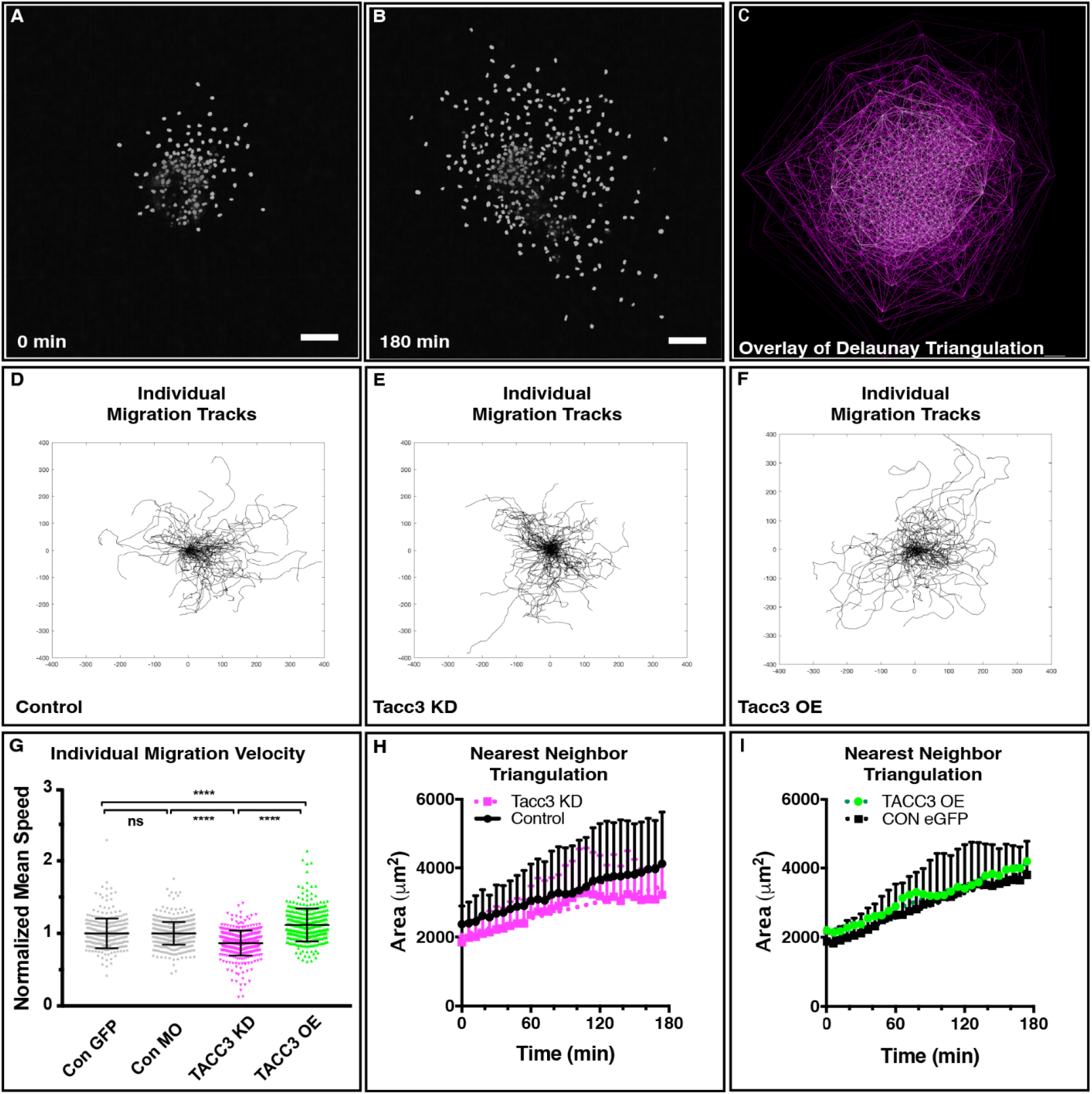
Tacc3 manipulation impacts cell velocity and dispersion dynamics of CNC cells. A-B) H2B-labeled control CNC cells undergoing migration away from the explant at 0 min and 3 hours post-migration. C) An overlay of Delaunay Triangulation demonstrates that both explant spread and the mean distance between nearest neighbors substantially increase over the course of migration. D-F) Representative cell tracks from all conditions (reoriented to start at a (0,0) origin) demonstrate generally low directional persistence in all conditions. G) Mean cell velocities are reduced in Tacc3 KD, while Tacc3 OE demonstrates increased velocity compared to controls, ^****^p ≤ 0.0001. H-I) Tacc3 KD results in reduced rates of ensemble dispersion over time, as measured by the change in area between nearest neighbors, while Tacc3 OE did not significantly affect ensemble dispersion. Scale bar 100 um.

A modest depletion in Tacc3 levels (∼20%) was sufficient to reduce mean cell velocity in individual CNC cells (Con: 1.82 ± 0.03 µm/min, n=351 cells, Tacc3 knockdown (KD): 1.51 ± 0.035 µm /min, n=334. p ≤ 0.0001, **Figure 1G**). This is the first demonstration that Tacc3 disruption can impact motility of primary, non-neuronal embryonic cells, and it supports a role for Tacc3 in maintaining normal CNC cell migration *in vivo*. This data is consistent with previous reports that TACC3 depletion in RPE1 cells could reduce cell velocity (Gutierrez-Caballero, Burgess et al. 2015). Similarly, TACC3 expression levels are dysregulated in most metastatic cancers, and increases in TACC3 expression correlate with invasive metastatic potential (Ha, Kim et al. 2013).

Thus, we next tested whether Tacc3 over-expression (OE) could increase individual CNC cell velocities. Indeed, a 20% percent increase in velocity of individual CNCs was noted with Tacc3 OE (Con: 1.75 ± 0.04 µm /min n=275, OE: 2.14 ± 0.03 µm /min, n=401, **Figure 1G**). No obvious defects in cell proliferation occurred at low-levels of Tacc3 KD (data not shown). This indicated that the reductions in total area occupied by CNC cells in Tacc3 KD embryos were likely due to reduced migration rates. However, it did not explain why these differences manifested as shorter and condensed CNC cell patches that still migrated effectively, albeit more slowly. We hypothesized that this may stem from changes in cell-cell interactions of individual CNC cells, which might then lead to ineffective ensemble dispersion.

To explore this, we performed measurements of total explant dispersion over time. Cells began tightly clustered within the explant and their migration led to rapid dispersion (**Figure 1A-B**). The area occupied by all explant cells typically expanded 2.5 – 4-fold over the course of 3 h. We did not find significant differences in total explant spread in either Tacc3 KD or OE, once data was normalized to initial explant size. However, this measure relies on the dynamics of cells on the periphery of the explant, and it is highly biased by the directional persistence of the cells. Thus, Delaunay triangulations were used to assess the mean distances between all neighboring cells within the explant (**Figure 1C**). A mean expansion of the triangle formed between nearest-neighbors over time would demonstrate effective ensemble dispersion (Carmona-Fontaine, Matthews et al. 2008; Szabó, Melchionda et al. 2016).

Tacc3-depleted cells showed a reduced rate of cell dispersion (**Figure 1H**). Initial triangle sizes were lower, indicating tightly-clustered explants, consistent with the slower rates of individual cell migration. Moreover, initial rates of dispersion were similar to that of controls. However, these rates plateaued over the course of migration, leading to an overall reduction in dispersion rate. This could indicate that Tacc3-depleted cells were unable to respond and repolarize effectively upon interaction with neighboring CNC cells. This would likely be less evident initially, as the tightly-clustered explant (0 min) would bias and facilitate effective dispersion. Tacc3 OE did not significantly affect total cell-cell ensemble dispersion (**Figure 1I**).

It has been shown that CNC cell motility *in vivo* is driven by transient attraction/repulsion dynamics between neighboring CNC cells (Carmona-Fontaine, Matthews et al. 2008; Woods, Carmona-Fontaine et al. 2014; Szabó, Melchionda et al. 2016). In the pharyngeal arches, CNC cells are directed ventrally by the chemoattractant Sdf1. However, rather than all cells moving persistently towards Sdf1, cells are funneled in the general direction of the cue, but are constantly undergoing cell-cell interactions with neighboring CNCs, by a balance of co-attraction and contact inhibition of locomotion (CIL). Direct cell contact drives CIL, in which a complete repolarization of directional migration pushes the colliding cells in opposite directions. Compared to a straightforward directional persistence towards a chemoattractant, this balance of directed migration, co-attraction, and cell dispersion was demonstrated to produce more efficient migration down the pharyngeal arches, when computationally modeled (Szabó, Melchionda et al. 2016). Thus, cell-cell ensemble dynamics are an important modulator of CNC cell behavior. Importantly, this behavior requires reconstruction of a new leading edge (Kadir, Astin et al. 2011). CIL is known to be driven by complete depolarization and repolarization of leading edge MT networks, and stabilization of a new leading edge is dependent on MT anchoring and stability at the cell cortex (Gundersen and Bulinski 1988). Therefore, Tacc3 manipulation could either impact cell ensemble behavior downstream of changes in MT stability or through changes on MT polymerization velocities.

### Tacc3 does not drive changes in MT stabilization

We first sought to establish whether Tacc3 could affect MT stability. One proxy for MT stabilization is the accumulation of the detyrosination post-translational modification of alpha-tubulin over time (Gundersen and Bulinski 1988; Morris, Nader et al. 2014; Pongrakhananon, Saito et al. 2018). However, to our surprise, under our culture conditions, CNC cells demonstrated essentially no alpha-tubulin detyrosination (**Figure 2A**). Conversely, all alpha-tubulin incorporated into the MT lattice was labeled positively for tyrosination, a marker of dynamic MT populations (**Figure 2B**). As it is not possible to ascertain changes in a non-existent signal, we needed to find another method to assess MT stability.

**Figure 2:**
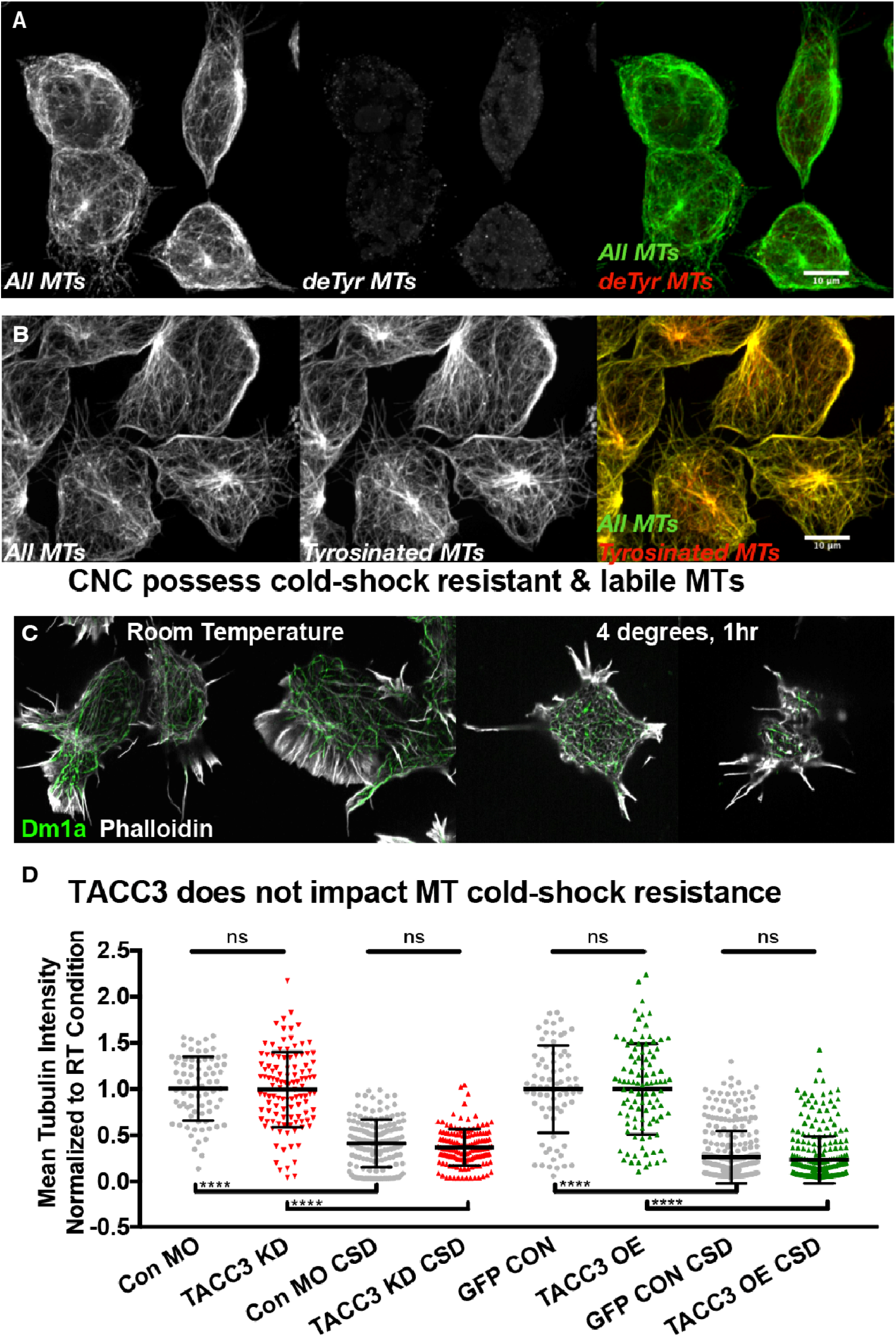
Tacc3 does not drive changes in MT stabilization. A-B) Post-translational modifications typically associated with MT stability were absent in CNC cells (A), but MTs stained positively for tyrosination, a marker of MT dynamicity (B). C) Despite lacking post-translation modifications associated with MT stability, cells treated with an extended cold-shock treatment had populations of cold-shock resistant MTs. C) Tacc3 KD and OE did not affect relative fluorescent intensity of remaining MTs following cold-shock treatment. Con MO, control condition for TACC3 KD; CSD, cold shock depolymerization. ^****^p ≤ 0.0001, Scale bar 10 um.

We next chose to examine MT network resistance-to-depolymerization as a proxy for MT stabilization, using thermodynamic MT disassembly (cold-shock depolymerization, or CSD) (Terasaki, Chen et al. 1986). Following one hr at 4°C, most MTs in CNC cells depolymerized, leaving a short astral array around the centrosome and a few fragments potentially associated with acentrosomal Golgi networks. (Given that *Xenopus laevis* cells are cultured at room temperature and without supplemented CO_2_, it is likely that amphibian cells possess modulations of the MT lattice that make them more resilient to CSD, as an hour at 4°C would likely gravely impact mammalian cells.) When cells were returned to room temperature, the MT network recovered rapidly, within ∼90 seconds (**Figure S2A**,**B**). Importantly, this occurred faster in Tacc3 OE cells, consistent with our observations that Tacc3 OE could drive increased MT growth velocities (**Figure S2C**) (Nwagbara, Faris et al. 2014). To address MT stability, we fixed and immunostained CNC cells with Dm1α, a pan-MT label, directly following their cold-shock treatment, without allowing MT network recovery (**Figure 2C**). Notably, CNC cells maintained 25-30% of their initial tubulin fluorescence, indicating that a subset of MTs maintained a resistance to CSD, even in the absence of post-translational modifications associated with MT stability. However, neither Tacc3 KD nor OE was sufficient to alter the proportion of MT fluorescent intensity that remained after cold-shock treatment (**Figure 2D**). Thus, our data does not support a role for Tacc3 in modulation of MT stability, which is consistent with previous data from neuronal growth cones (Erdogan, Cammarata et al. 2017).

### Tacc3 OE drives enhanced MT plus-end velocities in the leading edge

Given that MT network recovery in Tacc3 OE cells was faster following cold-shock treatment, we next questioned whether Tacc3 might impact cell motility by affecting leading edge MT dynamics. We previously reported that Tacc3 OE increased whole-cell MT plus end velocities by ∼15% in multiple embryonic cell types, including CNC cells (Nwagbara, Faris et al. 2014). Here, we repeated these assays, this time using CNC cells that expressed mKate2-MACF43, a minimal EB1-dependent +TIP (Honnappa, Gouveia et al. 2009). Using automated tracking of MT plus-ends, we observed that Tacc3 OE drove increased whole-cell MT plus-end velocities, consistent with our previous work (**Figure 3A**,**B**). We then used this data to perform a regional segmentation of local plus-end velocities. It is well established that MT growth velocities are faster in the cell central domain compared to its periphery (Wadsworth 1999; Komarova, Vorobjev et al. 2002; Alieva, Zemskov et al. 2010). Our results corroborated this, as central domain growth velocities in control CNC cells were statistically increased compared to those at the leading edge (**Figure 3C**). However, leading edge MT plus-end velocities in Tacc3 OE cells were not statistically different than those at the central domain. This indicates that while Tacc3 can affect plus-end growth velocities globally, its impact may be differentially enhanced at the leading edge of motile cells.

**Figure 3:**
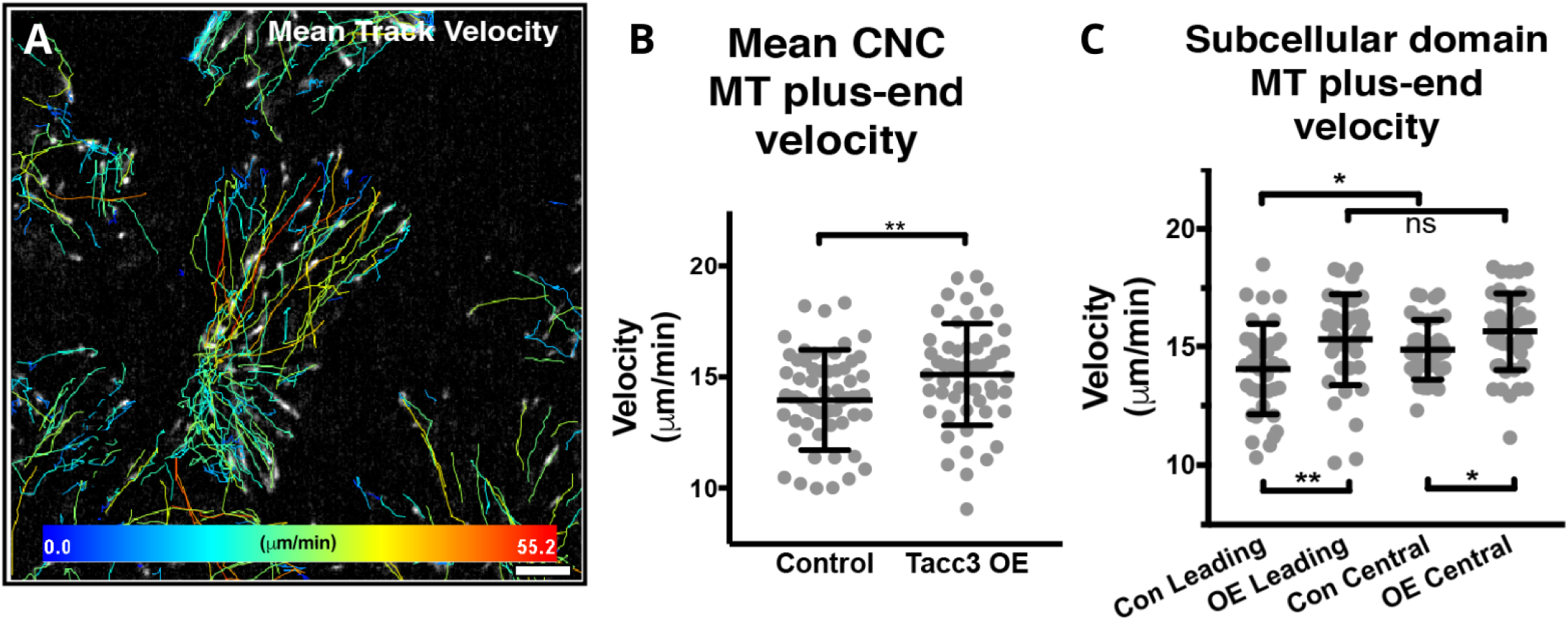
Tacc3 OE drives enhanced MT plus-end velocities in the leading edge. A) MT plus-end tracks color coded to designate velocity. B) Mean MT plus-end velocities in CNC cells are increased following Tacc3 OE. C) In control cells, mean MT plus-end velocities are enhanced in the central domain, while slow MT plus-end growth is shown in distal leading edges. However, Tacc3 OE enhances plus-end velocities in the leading edge, compared to controls. ^*^p ≤ 0.05, ^**^p ≤ 0.01, ns not significant.

### Tacc3 drives changes in focal adhesion disassembly

It remains unclear how Tacc3’s impact on MT growth velocities might drive changes in cell velocity. MT networks have occasionally been implicated in contributing to leading edge protrusion forces, but this process is generally accepted to be rate-limited by branched actin network polymerization dynamics (Pollard and Borisy 2003). Additionally, some of the established roles for MTs in modulating directed cell motility, such as establishment of polarity and directional persistence (Gundersen, Wen et al. 2005; Brodsky, Burakov et al. 2007), are associated with their regional stabilization, which we could not confirm downstream of Tacc3 manipulation. However, it is known that focal adhesion (FA) disassembly requires initial MT-FA targeting by dynamic MTs (Ezratty, Partridge et al. 2005; Broussard, Webb et al. 2008). Thus, we sought to examine whether Tacc3 could affect rates of FA remodeling (**Figure 4A**).

**Figure 4:**
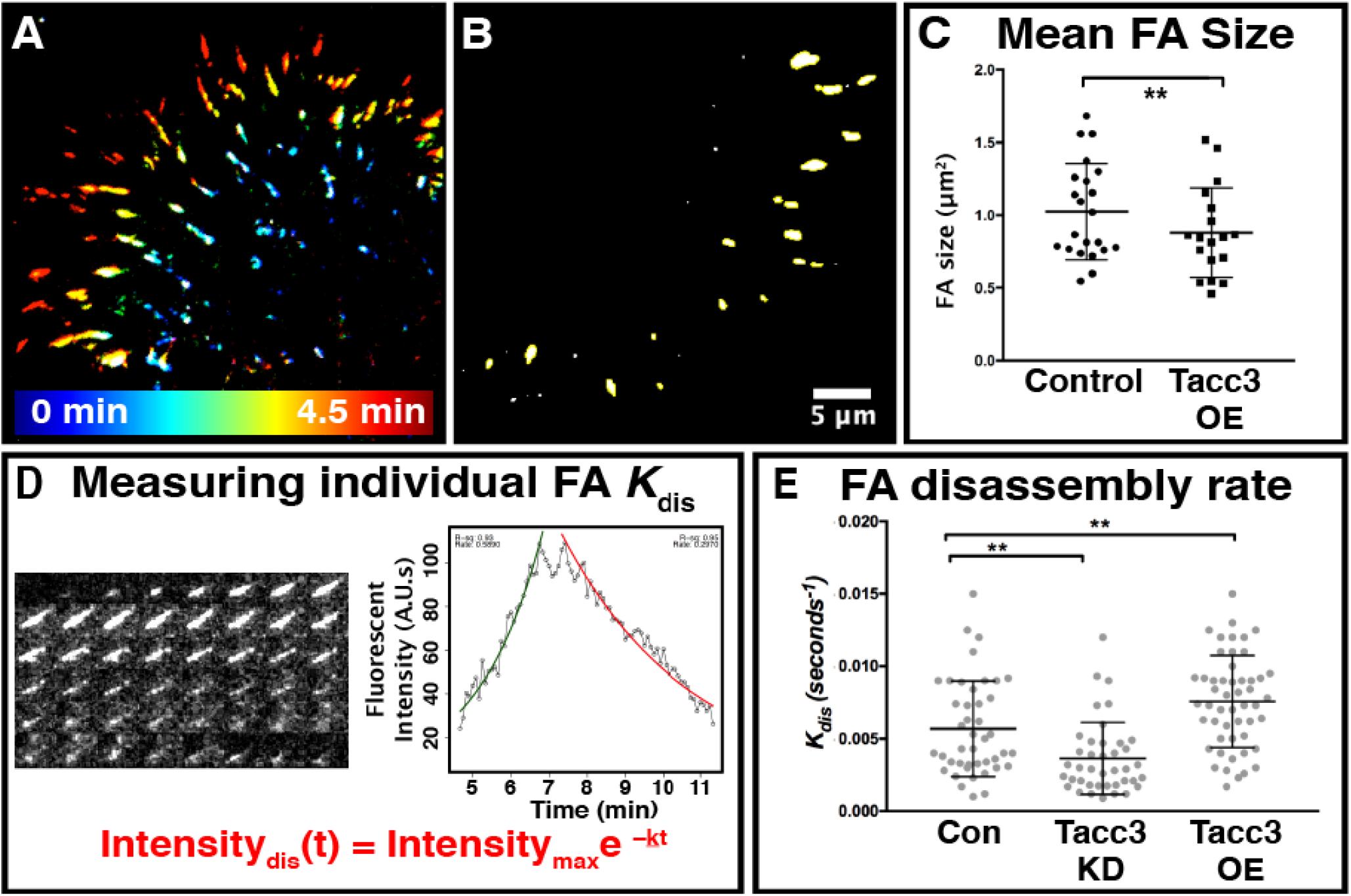
Tacc3 drives changes in focal adhesion disassembly. A) FAs in control CNC cells demonstrate very rapid turnover. B) Control FA size. C) Tacc3 OE reduces overall FA size, with no effect on FA number. D) The size-independent rate constant for FA disassembly (k_dis_) can be obtained by fitting the fluorescent intensity change during FA remodeling to a first-order exponential decay. E) CNC FA remodeling rates following Tacc3 KD and OE. Scale bar 5 um.

To test this, eGFP-Paxillin was co-expressed at low-levels, alongside Tacc3 manipulations, and imaged using total internal reflectance microscopy (TIRF), to observe impacts of Tacc3 OE on FA area and number. Mean FA size was reduced by Tacc3 OE, without any effects on total number (**Figure 4B**,**C**). However, the effect of FA size on cell velocity is controversial and often substrate-dependent (Gupton and Waterman-Storer 2006), and reduced FA size may be an indirect effect of cells that are already moving at any increased velocity. Thus, we next sought to describe rates of FA disassembly. CNC cell FAs demonstrated rapid assembly and turnover, with either phase generally lasting between 1-7 minutes. These rates are substantially faster than those documented in most epithelial wound or scratch assays, and necessitated rapid imaging intervals (at shorter durations) for accurate dynamics assessments. Fluorescently-labeled FA protein dissociation dynamics can be fit to a first-order exponential decay to obtain a rate of FA disassembly (**Figure 4D**) (Yadav and Linstedt). Importantly, this allows comparison of a size-independent rate constant, k_dis_, which is valuable given that we can assume that the smaller FAs present in Tacc3 OE cells would inherently disassemble at more rapid rates. Our data indicated that both Tacc3 OE and KD could indeed impact the rate of FA disassembly of CNC cells migrating on fibronectin (**Figure 4E**). As cell migration under these culture conditions is only persistent for transient bursts (such as over the course of the 90-second time-lapse used to measure MT dynamics in **Figure 3**), leading edge FAs rarely had the opportunity to grow, stabilize, and disassemble at a parallel lagging edge of the cell. Instead, CNC cells would frequently appear to ‘shift their weight’ sideways or backwards onto new or smaller eGFP-paxillin contacts to redirect motility. Given the regional enhancement of dynamic MT plus-end growth at the leading edge, these data suggest a model whereby Tacc3’s ability to mediate bulk- and leading edge MT plus-end velocities may facilitate rapid remodeling of transient, non-polarized contacts.

Altogether, our data suggests that Tacc3 may facilitate resilient MT network plasticity (**Figure 5**). Following CSD, CNC cells with Tacc3 OE demonstrated increased MT plus-end growth (**Figure S2**), whereas Tacc3 reduction decreased ensemble cell-cell dispersion (**Figure 1H**). Given that CIL is a primary driver of CNC cell dispersion, and as it necessitates a complete disassembly and repolarization of new leading edge MT arrays upon cell-cell contact, deficits in dispersion following Tacc3 KD could be attributed to deficits in MT regrowth. In support of this, we demonstrated regionally enhanced MT plus-end velocities at the leading edge of motile cells (**Figure 3C**). We also note that Tacc3 manipulation enhances rates of FA network remodeling (**Figure 4**). As CNC cells in our culture conditions demonstrate only transient polarity, FA disassembly was rarely persistently localized to one quadrant of the cell. In cells undergoing a chaotic, ‘drunk-walk’ behavior, dynamic remodeling of FAs may be enhanced when the MT network is plastic or perhaps even unstable.

**Figure 5:**
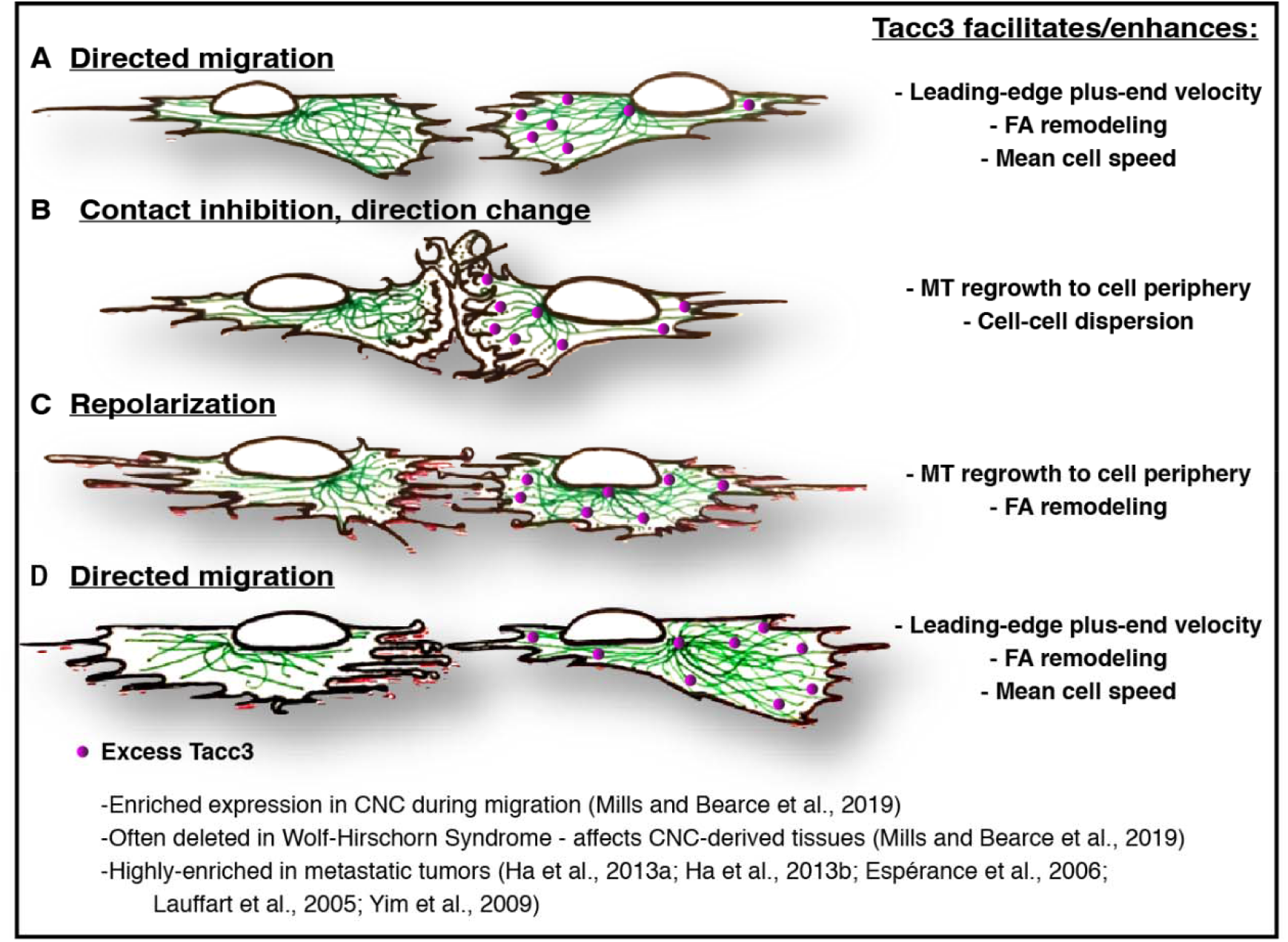
Tacc3 enhances cell velocity and dispersion. Tacc3 facilitates cell motility by way of enhancing MT dynamics and network regrowth. A) Two cells exhibiting directed motion. B-D) Following collision or directional change, Tacc3 facilitates MT regrowth to the cell periphery, demonstrates spatially-enhanced effects on leading-edge MT dynamics, and accelerates FA remodeling, to facilitate cell velocity and dispersion. (MTs = green; Adhesions = red/pink; Tacc3 = magenta)

Tacc3 is highly dysregulated in nearly all metastatic cancers, and the extent of this dysregulation has corresponded to an increase in metastatic potential (Lauffart, Vaughan et al. 2005; L’ Espérance, Popa et al. 2006; Yim, Tong et al. 2009; Ha, Kim et al. 2013; Ha, Park et al. 2013; Li, Ye et al. 2017; Matsuda, Miyoshi et al. 2018; Song, Liu et al. 2018; Qie, Wang et al. 2020). Thus far, studies of Tacc3 OE in tumors have not provided a clear mechanical model for these outcomes. However, our data may indicate an important link. Computational models have predicted that random migration patterns are advantageous to metastasizing cancer (Shatkin, Yeoman et al. 2020). While randomly-motile cells will take longer to reach any target, their probability of exploring any one region is enhanced. This behavior makes “random-walkers” statistically more likely to hit a ‘rare target’ in their environment. During tumor metastasis, these ‘rare targets’ could be transient weaknesses or openings in basement membranes. Our data does not inherently suggest that Tacc3 manipulation itself drives direct changes in the cell’s likelihood of adopting a random walk. Cells from control, KD, and OE conditions all demonstrated relatively similar oscillations between persistent and random motility. Further, Tacc3 manipulation showed no impacts on MT stabilization, which is frequently a mediator of directional persistence. Instead, our data suggest that Tacc3’s impact on CNC cell migration and tumor metastasis may derive from its ability to regulate whole-cell MT network plasticity.

## Materials and Methods

### Xenopus husbandry

Eggs obtained from female *Xenopus laevis* were fertilized *in vitro*, dejellied and cultured at 13-22°C in 0.1X Marc’s modified Ringer’s (MMR) using standard methods (Sive et al., 2010; Lowery et al., 2012). Embryos received injections of exogenous mRNAs or antisense oligonucleotide strategies at the two- or four-cell stage, using four total injections in 0.1X MMR containing 5% Ficoll. Embryos were staged according to Nieuwkoop and Faber (Faber and Nieuwkoop 1994). All experiments were approved by the Boston College and Boston University Institutional Animal Care and Use Committees and were performed according to national regulatory standards.

### Gene manipulation strategies

Capped mRNAs were transcribed and purified as previously described (Lowery et al., 2013; Nwagbara et al., 2014). Morpholino antisense oligonucleotides were used to target Tacc3 (5-AGTTGTAGGCTCATTCTAAACAGGA3), used and validated previously (Nwagbara, Faris et al. 2014). Morpholino KD conditions were always compared to cells from embryos injected with standard control morpholino (5-CCTCTTACCTCAGTTACAATTTATA-3) (Gene Tools, LLC). Tacc3 depletion was validated by Western Blot, as previously described (Nwagbara, Faris et al. 2014). Band detection was by chemiluminescence using Amersham ECL Western blot reagent (GE Healthcare BioSciences), and relative depletion was quantified by band densitometry using ImageJ. Rescue experiments were performed with exogenous mRNAs co-injected with their corresponding MOs. Plasmid for TACC3, cloned into pET30a, was a kind gift from the Richter lab (University of Massachusetts Medical School, Worcester, MA), which was subcloned into pCS2. As a start-site morpholino was utilized to block Tacc3 translation, a morpholino-resistant exogenous mRNA was generated by creating conserved mutations in the first 7 codons.

### Pharyngeal arch morphology quantification

Analysis of pharyngeal arches from *in situ* experiments was performed on lateral images in ImageJ, as previously described (Mills, Bearce et al. 2019). Measurements were taken to acquire: 1) Arch Area: The area of individual PA determined using the polygon tool, and 2) Arch Length: The length of the distance between the top and bottom of each PA. Statistical significance was determined using a student’s paired t-test in Graphpad (Prism).

### Neural crest explant isolation

A very helpful guide to neural crest isolation has been described previously (Milet and Monsoro-Burq 2014; Erdogan, Bearce et al. 2020). We offer only minor modifications here. Exogenous mRNAs and depletion strategies were injected as described above, and embryos were allowed to mature to Stage 18. Embryos were placed in modified DFA solution (53mM NaCl, 11.7 mM Na2CO3, 4.25 mM K Gluc, 2mM MgSO4, 1mM CaCl2, 17.5 mM Bicine, with 50ug/mL Gentamycin Sulfate, pH 8.3), before being stripped of vitelline membranes and imbedded in clay with the anterior dorsal regions exposed. Skin was removed above the neural crest using an eyelash knife, and CNC tissue was excised. Explants were plated on fibronectin-coated coverslips in imaging chambers filled with fresh DFA. Tissues were allowed to adhere, before being moved to the microscope for time lapse imaging of CNC cell motility.

### Cell Migration Assays

*Xenopus laevis* embryos were injected with 300pg of H2B-eGFP mRNA at the 4-cell stage, and CNC cell explants were excised and plated as above. pCS-H2B-EGFP was a gift from Sean Megason (Addgene plasmid 53744; RRID:Addgene 53744) (Megason 2009). CNC cell migration rates and dispersion assays were performed on a Zeiss Axio Observer inverted motorized microscope, using a Zeiss AxioCam camera controlled with Zen software. Images were captured at 20x (DIC and GFP), using 4×4 tiled acquisitions to capture the entire migratory field. Eight to ten explants, from both control and experimental conditions were imaged, at a six-minute interval, for three hours. Data was imported to Fiji, background subtracted, and cropped to a uniform field size for the analyses below.

### Cell Speed Assays

Migration tracks of individual cells were collected using automated tracking (Trackmate) (Tinevez, Perry et al. 2017). Tracks were filtered by length, to use only tracks that contained at least 10 frames of data, and quality. The highest quality 25% of the remaining tracks were used for velocity measurements. This removed tracks from the inside of the explant, where track accuracy was difficult to manually verify due to the close proximity of nearest neighbors. Average final number of tracks collected per tissue explant was between 100-250. Mean velocities of each track (per cell) were imported to Prism (Graphpad), and compared between conditions using unpaired t-tests. Three independent experiments were performed for each experimental condition.

### Directionality Assays

For directionality and persistence visualization, 100 tracks from the filtered cell speed assays above were randomly withdrawn from each condition (randomization performed with matrix manipulation, MATLAB), and plotted from a (0,0) origin. Displacement, Directionality, and RandomWalk Analysis were calculated from the individual x,y coordinate positions in the Spots in Tracks file.

### Explant Dispersion

Explant dispersion analysis was performed using the same migration assay data sets gathered for cell speed and directionality analysis, but the experiments were re-tracked (Trackmate) (Tinevez, Perry et al. 2017) to detect all nuclei in each frame, with no regard for track accuracy. Raw x,y coordinates of each particle (nuclei) were extracted to a new array of ROIs (each x,y coordinate as a single center of mass), sorted by their corresponding frame, and imported to the ROI Manager. A looping command was used to generate Delaunay Voronoi Triangulations for each individual frame, using the Delaunay Voronoi plug-in (FiJi). Triangulations from each frame were concatenated to create dispersion movies. To get mean triangle and explant dispersion measurements, Analyze Particles was used to count and measure triangle area. Triangle areas for each frame were summed to generate total explant area. Plots of total explant dispersion (sum area of all triangles) were normalized to initial area. Average areas of individual triangles were plotted as non-normalized raw values. Rates of growth were compared between experimental conditions with student t-tests in Prism (Graphpad).

### MT dynamics Assays

To measure MT growth velocities, embryos were injected with 100 pg/embryo of mKate2-MACF mRNA, prior to neural crest dissection as describe above. Images of mKate2-MACF labeled plus-ends from individual cells were collected over a 1.5 minute duration, at a 2 second imaging interval. Comet tracking was performed using plusTipTracker (Applegate, Besson et al. 2011; Lowery, Stout et al. 2013). We previously validated the imaging conditions and tracking parameters used in this study for accurate detection of EB1 comets in *Xenopus laevis* growth cones (Stout, D’Amico et al. 2014), and in neural crest (Nwagbara, Faris et al. 2014). The same parameters were used for all movies: maximum gap length, eight frames; minimum track length, three frames; search radius range, 512 pixels; maximum forward angle, 50°, maximum backward angle, 10°; maximum shrinkage factor, 0.8; fluctuation radius, 2.5 pixels; time interval, 2 s. Average MT growth speed, lifetime, and lengths were pulled ROIs within either the leading and central domains; regions that did not contain at least 10 MT comets were discarded. Because MT dynamics parameters were compiled from multiple individual experiments, and there can be significant day-to-day fluctuations in control MT dynamics, the final compiled data were normalized relative to the mean of the control data for each experiment.

### FA Remodeling

Live imaging of FA remodeling was performed in CNCs from embryos injected with eGFP-Paxillin (100 pg/embryo). pCS2-Paxillin-eGFP was a gift from Tim Gomez (U of Wisconsin). Total internal reflection fluorescence (TIRF) imaging was performed using an inverted Nikon Ti-E system, with a 60x Plan-Apo 1.49 lens and an Andor Zyla sCMOS camera. Images were collected every 5 seconds for 15-20 minutes, with a 100 ms exposure without binning, at 15% laser power. FA remodeling was analyzed as discussed previously (Stehbens and Wittmann 2014); briefly, fluorescence intensities of individual FAs were background subtracted and plotted over time, using a 10-frame median window for smoothing. A first order exponential decay was fitted to the disassembly phase, and k_*dis*_ was used as a size-independent metric of FA disassembly.

### Immunostaining

For assessment of MT stability following Tacc3 manipulation, immunocytochemistry was performed to label post-translational modifications associated with stable or dynamic MT populations, and the area of labeling was compared to that of a global MT label. Neural crest explants were dissected, and allowed to migrate 4 hours prior to fixation. Cultures were fixed for 10 minutes in 0.2% glutaraldehyde and 0.1% Triton X-100 in PHEM buffer, as previously described (Challacombe, Snow et al. 1997). Sodium borohydride quenching solution (1 mg/mL in PBS) was applied for 15 min. Cultures were blocked for one hour with 5% goat serum in PBS and 0.1% Triton X-100. MTs were labeled with rat monoclonal antibody YL 1/2 directed against tyrosinated -tubulin (1:1000, Millipore), or rabbit polyclonal antibody against detyrosinated -tubulin (1:1000, Abcam), to label dynamic or stable MT populations, respectively, in addition to mouse monoclonal alpha-tubulin (Dm1a) (1:1000, Abcam) to label all MTs. Primary antibodies were diluted into a blocking solution and then applied for 45 min. Coverslips were rinsed for 15 min in PBS, and incubated with corresponding fluorescent secondary antibodies applied in blocking solution for 1 hour (Goat Anti-Mouse, Goat Anti-Rabbit, or Goat Anti-Rat, conjugated to Alexa488 or Alexa568; 1:1000, Lifetechnologies). For cold-shock assays, only DM1a was used for MT labeling; an Alexa-488 conjugated Phalloidin (1:500, Lifetechnologies) was applied with secondary, to serve as a whole-cell area boundary marker. Cultures were rinsed for 15 minutes in PBS, and solutions were replaced with glycerol mounting media. Images were acquired within 10-12 hours of staining.

### Whole Mount in situ Hybridization

Embryos were fixed overnight at 4°C in a solution of 4% paraformaldehyde in phosphate-buffered saline (PBS), gradually dehydrated in ascending concentrations of methanol in PBS, and stored in methanol at -20°C for a minimum of two hours, before *in situ* hybridization, which was performed on fixed embryos as previously described (Saint-Jeannet 2017). After brief proteinase K treatment, embryos were bleached under a fluorescent light in 1.8X saline-sodium citrate, 1.5% H2O2, and 5% (vol/vol) formamide for 20 minutes to 1 hour before prehybridization. During hybridization, probe concentration was 0.5 ug/mL. The *tacc3* construct used for a hybridization probe was subcloned into the pGEM T-easy vector (Promega, Madison, WI). The Xenopus *twist* hybridization probe was a gift from Dr. Dominique Alfandari (University of Massachusetts at Amherst, MA), which was subcloned into the pCR 2.1TOPO vector (AddGene, Cambridge, MA). The antisense digoxigenin-labeled hybridization probes were transcribed in vitro using the T7 MAXIscript kit. Embryos were imaged using a Zeiss AxioCam MRc attached to a Zeiss SteREO Discovery.V8 light microscope.

## Acknowledgements

We thank members of the Lowery lab, especially Mackenzie Hulme, for helpful comments. We thank Nancy McGilloway and Todd Gaines for excellent *Xenopus* husbandry. We also thank the National Xenopus Resource (RRID:SCR-013731) and Xenbase (RRID:SCR-003280) for their invaluable support to the model organism community. This work was funded by the NIH National Institute of Dental and Craniofacial Research (R03 DE025824), the NIH National Institute of Mental Health (MH109651), the March of Dimes (1-FY16-220), Charles H. Hood Foundation 2018-2019 Bridge Funding Award from Charles H. Hood Foundation, Inc., Boston, MA, and the American Cancer Society (RSG-16–144-01-CSM). EB was funded by a National Science Foundation Graduate Research Fellowship.

## Abbreviations

MT: microtubule
CNC: cranial neural crest
FA: focal adhesion
KD: knockdown
OE: overexpression
CIL: contact inhibition of locomotion

## Supplementary Figures

**Figure S1.**
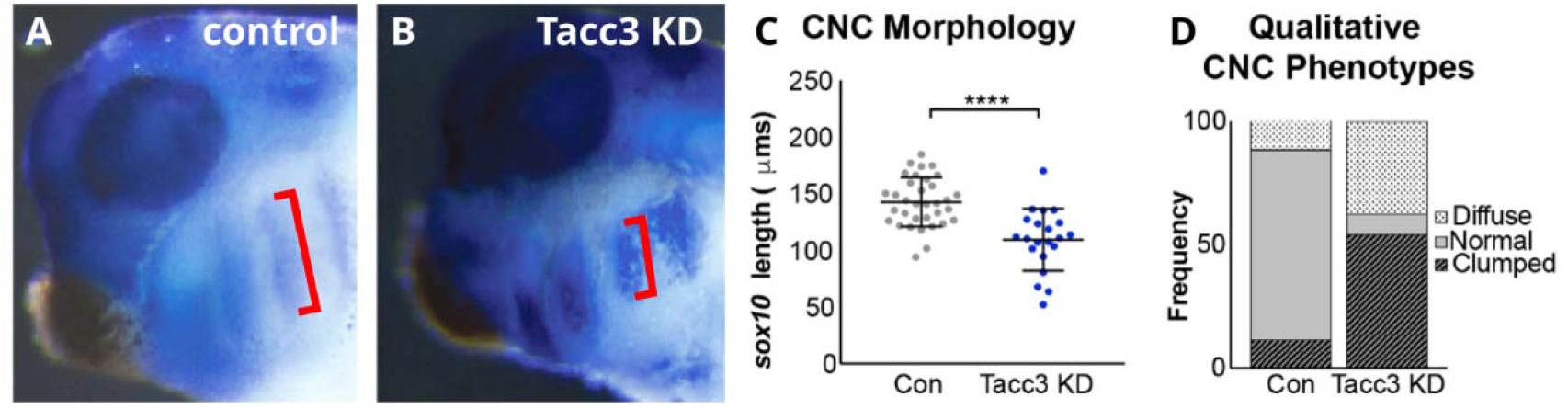
Tacc3 KD impacts CNC cell migration *in vivo*. (A,B) *Sox9 in situs* of stage 25 *Xenopus laevis* embryos. Left side view of embryo. Red bracket denotes pharyngeal arch length. (C,D) Quantitative and qualitative analysis of *sox9* expression.

**Figure S2.**
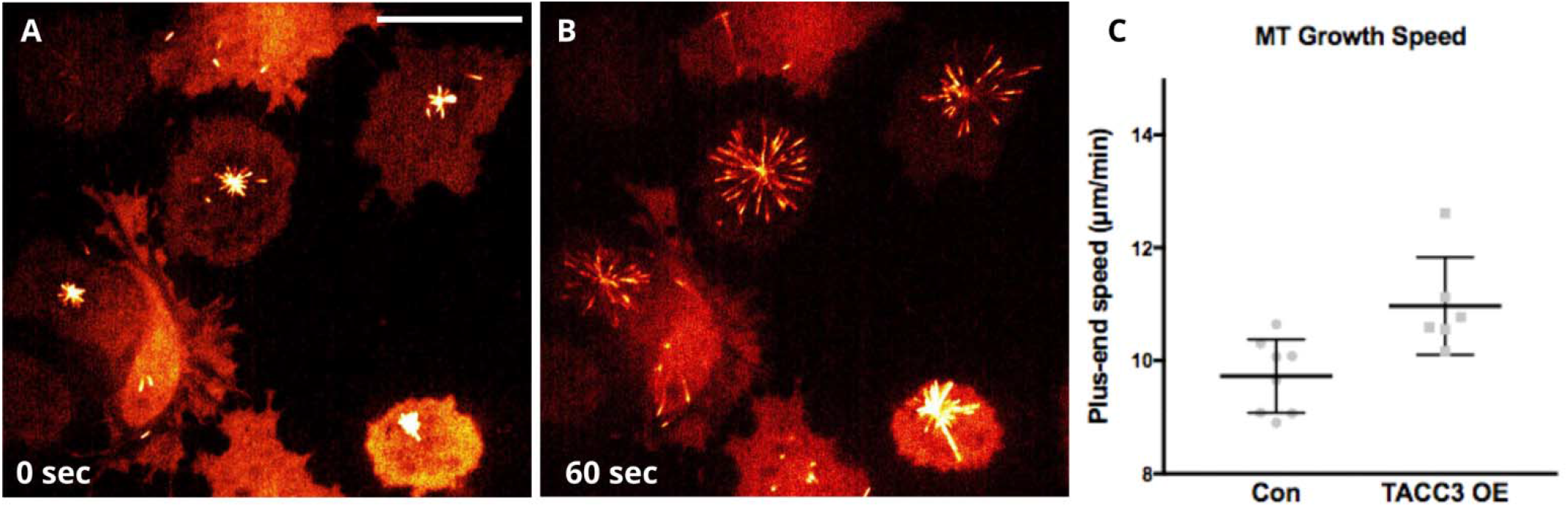
MT network recovery after cold-shock depolymerization. (A) Cells, expressing fluorescent plus-end tracking protein, immediately after 1 hour at 4 degrees Celsius. (B) 60 seconds after recovery. (C) MT plus-end growth velocities during recovery. Scale bar is 25 um.

